# *Mutator* transposon insertions within maize genes often provide a novel outward reading promoter

**DOI:** 10.1101/2023.06.05.543741

**Authors:** Erika L. Ellison, Peng Zhou, Peter Hermanson, Yi-Hsuan Chu, Andrew Read, Candice N. Hirsch, Erich Grotewold, Nathan M. Springer

**Affiliations:** Department of Plant and Microbial Biology, University of Minnesota, Saint Paul, MN 55108; Department of Biochemistry and Molecular Biology, Michigan State University, East Lansing, Michigan 48824; Department of Agronomy and Plant Genetics, University of Minnesota, Saint Paul, MN 55108

**Keywords:** Mutator, maize, transposable elements

## Abstract

The highly active family of *Mutator* (*Mu*) DNA transposons has been widely used for forward and reverse genetics in maize. There are examples of *Mu*-suppressible alleles which result in conditional phenotypic effects based on the activity of *Mu*. Phenotypes from these *Mu*- suppressible mutations are observed in *Mu*-active genetic backgrounds, but absent when *Mu* activity is lost. For some *Mu*-suppressible alleles, phenotypic suppression likely results from an outward-reading promoter within *Mu* that is only active when the autonomous *Mu* element is silenced or lost. We isolated 35 *Mu* alleles from the UniformMu population that represent insertions in 24 different genes. Most of these mutant alleles are due to insertions within gene coding sequences, but several 5’ UTR and intron insertions were included. RNA-seq and *de novo* transcript assembly were utilized to document the transcripts produced from 33 of these *Mu* insertion alleles. For 20 of the 33 alleles, there was evidence of transcripts initiating within the *Mu* sequence reading through the gene. This outward-reading promoter activity was detected in multiple types of *Mu* elements and doesn’t depend on the orientation of *Mu*. Expression analyses of *Mu*-initiated transcripts revealed the *Mu* promoter often provides gene expression levels and patterns that are similar to the wild-type gene. These results suggest the *Mu* promoter may represent a minimal promoter that can respond to gene *cis*-regulatory elements. Findings from this study have implications for maize researchers using the UniformMu population, and more broadly highlights a strategy for transposons to co-exist with their host.

**Article Summary:** *Mutator* (*Mu*) transposable elements are a widely used tool for insertional mutagenesis in maize and often insert in the 5’ regions of genes. The characterization of transcripts for *Mu* insertion alleles reveals complex transcripts. These often result in one transcript that covers the first portion of the gene terminating in *Mu* and a second transcript initiating within *Mu* covering the latter portion of the gene. This may reflect a strategy for *Mu* to minimize the consequences of insertions within genes.

## Introduction

Transposon insertion stocks have been developed and successfully used to study gene function in several organisms, including plants (Parinov and Sundaresan 2000; Brutnell 2002; Østergaard and Yanofsky 2004; Tadege *et al*. 2005), invertebrate animal models (Cooley *et al*. 1988; Bessereau *et al*. 2001; Thibault *et al*. 2004; Bessereau 2006), bacteria (Cain *et al*. 2020), and a variety of single-cell eukaryotes (Guo *et al*. 2013; Michel *et al*. 2017). Across species, the applicability of these transposon stocks vary due to properties of the transposon/transposase and the target genome, including sufficient transpositional activity, endogenous transposon copy number, transposon element type, family and size, integration site preference, and chromatin landscape of the genome (Ivics *et al*. 2009). In maize, there are two widely utilized DNA transposon families with sequence-indexed libraries: *Activator/Dissociation (Ac/Ds)* (Brutnell and Conrad 2003; Vollbrecht *et al*. 2010) and *Mutator (Mu)* (McCarty *et al*. 2005). Multiple *Mu* transposon populations have been generated in maize, including UniformMu (McCarty *et al*. 2005), BonnMu (Marcon *et al*. 2020), Mu-Illumina (Williams-Carrier *et al*. 2010), Pioneer Hi-Bred International’s Trait Utility System for Corn (TUSC) (Briggs and Meeley 1995; Bensen *et al*. 1995), and Maize-Targeted Mutagenesis population (MTM) (May *et al*. 2003). Both *Mu* and *Ds* elements preferentially transpose into low copy sequences or genic regions which is useful for mutagenesis (Bennetzen and Springer 1994; Rabinowicz *et al*. 1999; Hanley *et al*. 2000; Fernandes *et al*. 2004; McCarty *et al*. 2005). The utility of *Ac/Ds* stocks for mutagenesis is somewhat limited due to low copy number, low germinal insertion frequency, and the tendency for new copies to insert into sites genetically linked to the donor loci (Vollbrecht *et al*. 2010). By contrast, *Mu* stocks are more widely used for reverse genetics because *Mu* has a high germinal insertion frequency, high forward mutation rate, and frequently inserts into genic regions that are unlinked to the donor loci (Lisch *et al*. 1995; Cresse *et al*. 1995; Settles *et al*. 2004; Fernandes *et al*. 2004; McCarty *et al*. 2005; Vollbrecht *et al*. 2010).

*Mu* transposable elements are the most mutagenic plant transposons known, due to their high transposition frequency and tendency to insert into low copy sequences or genic regions (Cresse *et al*. 1995; Lisch 2002; Settles *et al*. 2004; Fernandes *et al*. 2004; McCarty *et al*. 2005). The original *Mu* element was first described in maize in 1978 by Donald Robertson (Robertson 1978, 1983). Robertson identified a maize line with what he called “mutator” activity that had a very high forward mutation rate, 50- to 100-fold greater than that of the background. Upon outcrossing mutator plants to non-mutator plants, 90% of the progeny retained the high mutation frequency, which suggested there is non-Mendelian inheritance of “mutator” activity. It is now known that *Mu* transposons were responsible for the mutations in Robertson’s lines (Strommer *et al*. 1982; Bennetzen 1984; Taylor and Walbot 1987; Fedoroff and Chandler 1994; Lisch and Jiang 2015). There is evidence that *Mu* elements in maize can reach high copy numbers because they can transpose at frequencies nearing 100% (Alleman and Freeling 1986; Walbot and Rudenko 2002), have a high germline mutation rate, and are rarely excised from the germline (excision rate < 10^-4^) (Schnable and Peterson 1988; Levy *et al*. 1989; Brown *et al*. 1989; Walbot 1991; Levy and Walbot 1991; Bennetzen 1996).

The *Mutator* system is a two-component system with one autonomous element, *MuDR*, and several non-autonomous elements (*Mu1* to *Mu13*) (Lisch 2002; Tan *et al*. 2011). Non-autonomous *Mu* elements all contain similar ∼220 bp terminal inverted repeats (TIRs), but each class of element has unique internal sequences (Chandler and Hardeman 1992; Lisch and Jiang 2015). *Mutator* activity is dependent on the presence of an active autonomous *MuDR* element to encode the proteins necessary for transposition of itself and non-autonomous elements (Chomet *et al*. 1991b; Hershberger *et al*. 1991). *Mu*, like *Ds*, preferentially inserts into gene rich regions; however, *Mu* exhibits a stronger preference than *Ds* for 5’ UTR or promoter regions (Dietrich *et al*. 2002; Vollbrecht *et al*. 2010). These genic regions where *Mu* lands have distinct chromatin, DNA methylation, and recombination activity that could influence *Mu* element targeting (Bennetzen 2000; Lisch and Jiang 2015). *Mutator* activity can be epigenetically regulated, such that some plants with *MuDR* are in fact *Mu*-inactive due to heterochromatin-mediated silencing of the *MuDR* coding sequences, which is accompanied by high levels of DNA methylation (Chandler and Walbot 1986; Chomet *et al*. 1991b; McCarty *et al*. 2005). In plants containing epigenetically silenced *MuDR* the non-autonomous *Mu* elements do not exhibit evidence for transposition and are methylated in the *Mu*-inactive state. In the *Mu*-active state, the autonomous and non-autonomous *Mu* elements are hypomethylated and products from *MuDR* can mobilize *Mu* elements (Lisch *et al*. 1995; Hershberger *et al*. 1995). To develop UniformMu populations in maize that are genetically stable (in *Mu*-inactive genetic backgrounds) new germinal *Mu* insertions from lines with *MuDR* activity (*Mu*-active) are stabilized by selecting against somatic transposition of *Mu* using the *bronze1-mum9* (*bz1*-*mum9*) mutation as a genetic marker for *MuDR* activity (Brown *et al*. 1989; Brown and Sundaresan 1992; McCarty *et al*. 2005, 2013; Settles *et al*. 2007).

*Mutator* has been used to mutagenize many genes and isolate loss-of-function alleles (Chen *et al*. 1987; Stinard *et al*. 1993; Greene *et al*. 1994; Dietrich *et al*. 2002; Bortiri *et al*. 2006; Settles *et al*. 2007). In many cases these *Mu* insertion alleles produce a stable phenotype that does not change depending upon the epigenetic state of *Mu*. However, there is evidence that the phenotypic consequences for some of these *Mu*-induced alleles can be suppressed depending on the state of *Mu*. *Mu*-induced mutations that are suppressible, *Mu*-suppressible alleles, exhibit a mutant phenotype in *Mu*-active genetic backgrounds that can be suppressed, returning to a wild-type phenotype, when *Mu* activity is lost (*Mu*-inactive). *Mu*-suppressible alleles were well characterized in a recessive loss-of-function mutation, *hcf106-mum1*, caused by insertion of *Mu1* in the promoter of *HCF106*, a gene in maize required for chloroplast membrane biogenesis (Martienssen *et al*. 1989, 1990; Barkan and Martienssen 1991). Homozygous *hcf106-mum1* maize seedlings expressed a non-photosynthetic, pale green mutant phenotype only in the absence of *Mu* activity (*Mu*-inactive) (Martienssen *et al*. 1989, 1990; Barkan and Martienssen 1991; Das and Martienssen 1995). It was found that in *Mu*-inactive stocks an outward-reading promoter near the termini of *Mu* can direct transcription outward into the adjacent gene and substitute for the *HCF106* promoter (Barkan and Martienssen 1991). Since this discovery several other Mu-suppressible alleles have been described, including *Les28* (Martienssen and Baron 1994), *a1-mum2* (Chomet *et al*. 1991b; Pooma *et al*. 2002), *rs1* and *lg3* (Girard and Freeling 2000), *kn1* (Lowe *et al*. 1992), and *rf2a* (Cui *et al*. 2003). The full extent and molecular mechanisms of suppressibility of *Mu*-induced mutations have not been characterized widely, but may be a property of the *Mutator* system (Lisch and Jiang 2015).

We sought to further characterize the frequency of promoter activity in *Mu*-induced mutations and properties influencing the promoter’s ability to direct transcription outward has not been previously reported. Here we characterized the transcripts of 33 *Mu* insertion alleles. We find evidence that many (n = 20) of these alleles result in the production of two transcripts: one initiating at the wild-type gene transcription start site (TSS) and another initiating from a *Mu* outward-reading promoter. This *Mu* outward-reading promoter appears to be functional in several of the non-autonomous *Mu* elements and is not dependent upon *Mu* orientation.

Interestingly, our findings suggest that the *Mu* promoter is a minimal promoter that often shows expression levels and patterns quite similar to the gene it is inserted within. These findings highlight a potential strategy for co-evolutionary interactions between transposons and their host genomes.

## Results

### Characterization of transcripts arising from genes with *Mutator* insertions

To investigate the effect of *Mutator (Mu)* insertions on transcript structure, we isolated homozygous mutants for 35 insertions in 24 genes (McCarty *et al*. 2005, 2013). These included 8 insertions in 5’ UTR sequence, 22 insertions in coding regions, and 5 insertions in introns (Figure 1, Table S1). These frequencies do not necessarily reflect the spectrum of insertion sites for all *Mu* elements. We focused on selection of insertions within coding sequences as the mutants were originally selected as putative loss-of-function alleles for maize transcription factors. We generated RNA-seq data for three biological replicates of each homozygous mutant and wild-type allele. A single tissue for each mutant allele was selected to generate RNA-seq data based on evidence of wild-type allele expression (Table S2). The expression level of the mutant allele was documented by aligning RNA-seq data to the W22 reference genome (Springer *et al*. 2018) and only 8/35 alleles exhibited significantly lower transcript abundance in the mutant relative to the wild-type (Table S1). The majority, 25/35, of mutant alleles do not have any significant change in transcript abundance relative to wild-type, and two alleles have significantly higher transcript abundance (Table S1). Assessment of the mapped transcript reads derived from homozygous mutant plants revealed reduced coverage at the site of the *Mu* insertion (Figure S1). The drop in coverage flanking the *Mu* insertion site is expected if there is a novel junction and/or sequence present at this region in the mutant allele transcript relative to the W22 reference genome.

**Figure 1.**
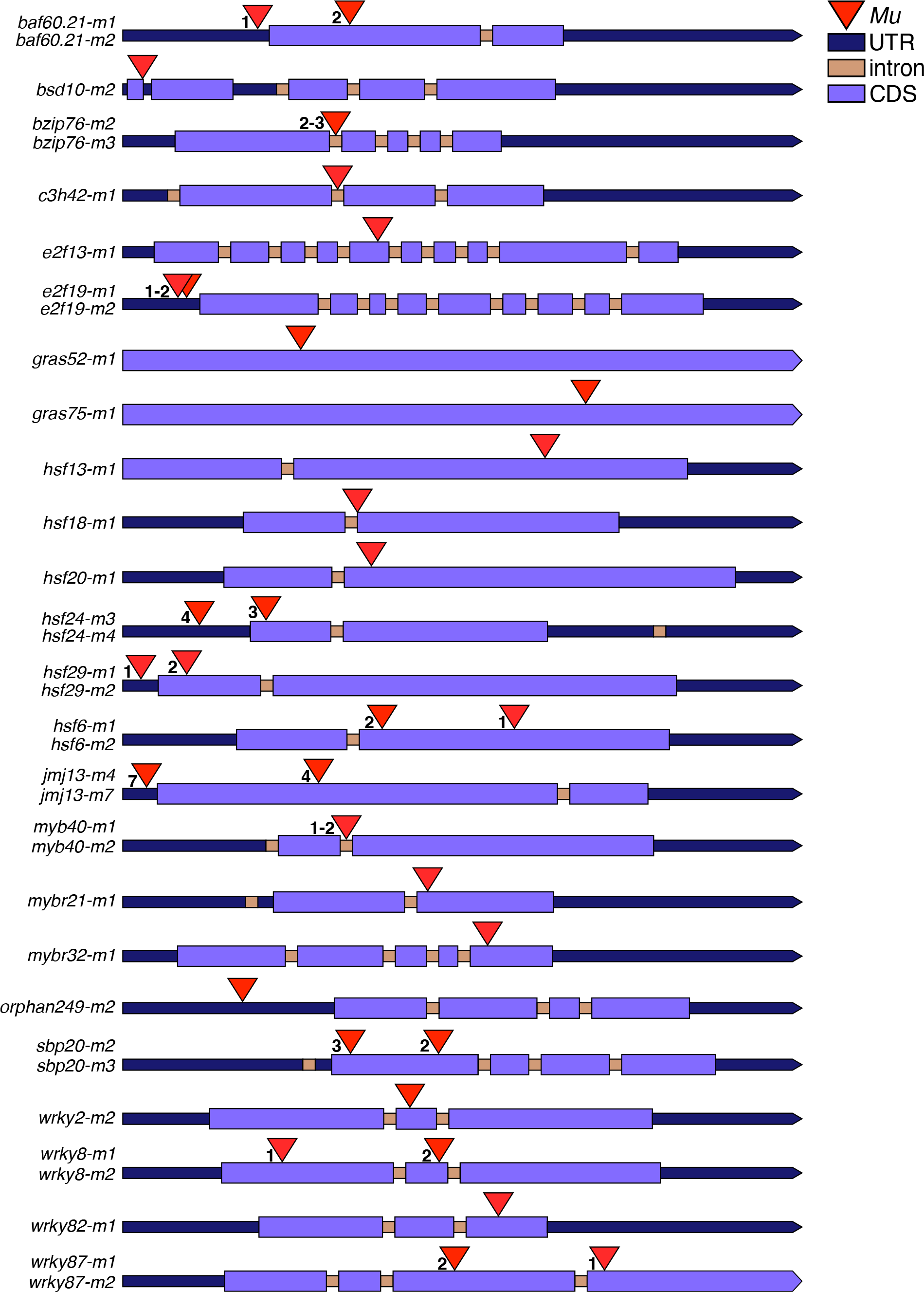
Schematic of *Mutator* insertion locations for 35 UniformMu mutants. The W22 gene models are indicated by different colors/shapes to represent UTRs, coding sequence and introns. The UTRs and CDS’ for each gene model are scaled proportionally, but introns are not to scale. *Mu* transposon insertions are indicated by red triangles and independent mutant alleles from the same gene are depicted by numbers. The *BSD10* (*Zm00004b040474*) gene model is based on the B73v4 (*Zm00001d026518*) gene annotation due to a fused gene annotation in W22.

There are several potential transcript structures that might be expected to be produced from *Mu* insertion alleles in comparison to the full-length wild-type W22 gene transcripts (Figure 2A). Mutant allele transcripts could include: read-through transcription resulting in retention of the full *Mu* sequence (*Mu* read-through transcript), novel splicing events that include retention of a portion of the *Mu* sequence (*Mu* spliced transcript), transcript initiation at the wild-type gene TSS with premature termination in the *Mu* sequence (gene TSS-*Mu* transcript), or transcript initiation from a *Mu* outward-reading promoter reading through to the wild-type termination site (*Mu* TSS transcript) (Figure 2A). These potential *Mu* insertion allele transcript structures are not necessarily mutually exclusive. To determine if *Mu* insertion alleles produced read-through transcripts with all or a portion of *Mu* sequence retained (*Mu* read-through or *Mu* spliced transcripts) we performed RT-PCR with gene-specific primers that flank the *Mu* insertion site (Figure 2B, Table S4). Although we were able to amplify the expected product in wild-type plants we were not able to detect amplified products in plants homozygous for the *Mu* insertion for 7 mutant alleles that were tested (see examples in Figure 2C). This suggests that read- through transcription of the *Mu* insertion with retention of partial or complete *Mu* sequences in the mRNA is unlikely or rare.

**Figure 2.**
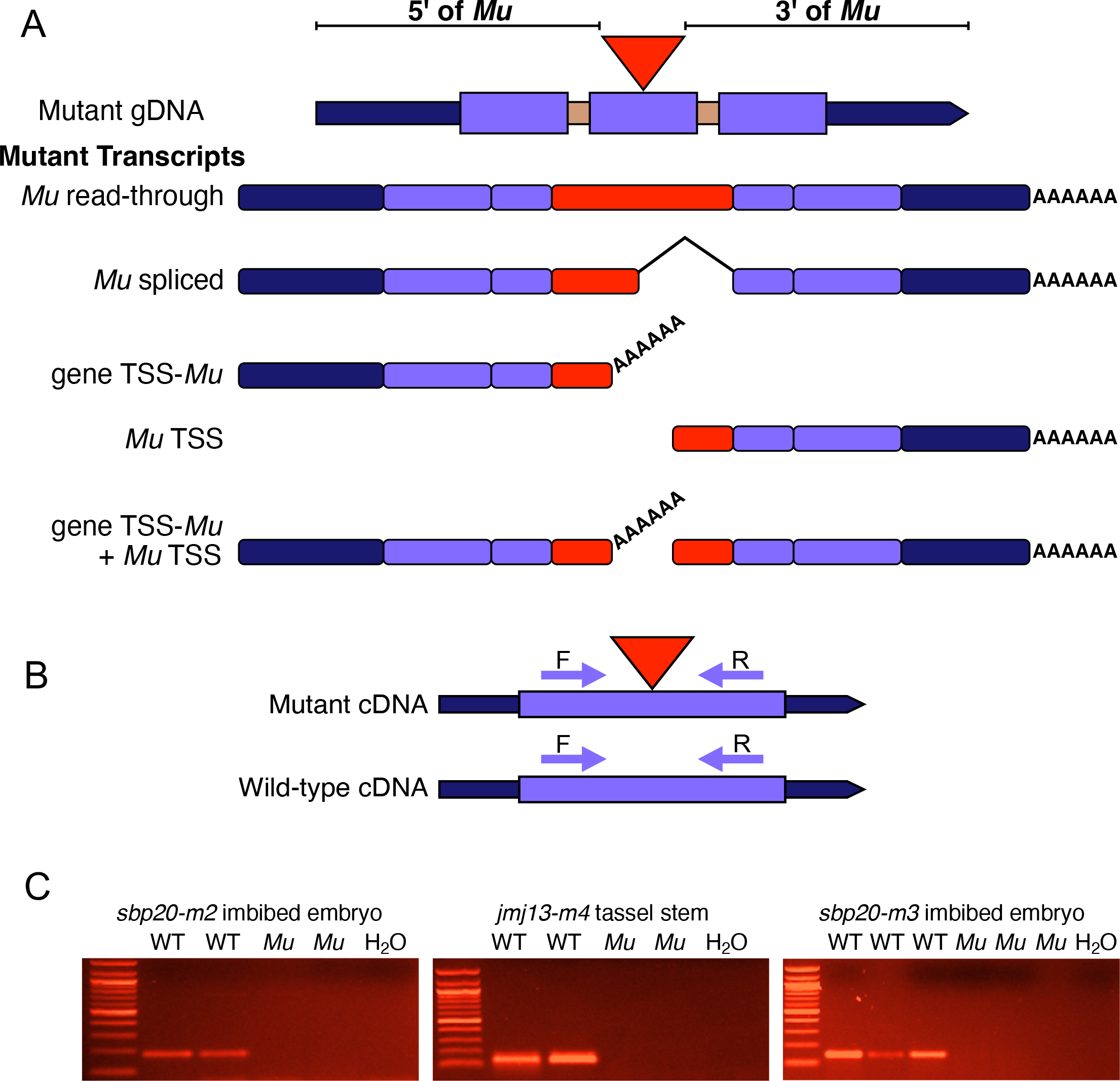
Potential *Mu* insertion allele transcript structures. A) Schematic of 5 mutant transcript structures that could result from a *Mu* transposon insertion. Potential transcripts include: a transcript with gene sequences 5’ and 3’ of the *Mu* insertion and all or a portion of *Mu* retained (*Mu* read-through), a transcript with sequence 5’ and 3’ of the *Mu* insertion and partial *Mu* sequence retained due to alternative splicing (*Mu* spliced), a partial transcript initiating at the gene TSS and terminating in *Mu* (gene TSS-*Mu*), a partial transcript initiating within *Mu* and reading through the 3’ gene sequence (*Mu* TSS), or both the gene TSS-*Mu* and *Mu* TSS transcripts. **B)** Schematic of RT-PCR primer design to test for *Mu* read-through and *Mu* spliced transcripts with gene specific primers (F and R) flanking the annotated *Mu* insertion site in mutant and wild-type alleles. A larger PCR product will be amplified in the mutant allele relative to wild-type if all or a portion of *Mu* is retained. No product will be amplified in the mutant if there are transcripts initiating or terminating in *Mu*. **C)** RT-PCR gels of gene-specific primers flanking *Mu* for at least two biological replicates of 3 mutant and wild-type alleles. All three alleles lack mutant cDNA amplification relative to wild-type.

To investigate transcript structure of the *Mu* insertion alleles, we generated *de novo* transcript assemblies for each mutant (35 alleles for 24 genes) and wild-type W22 RNA-seq (5 tissues) dataset for a total of 40 transcriptome assemblies. The *de novo* transcriptome assemblies for the W22 control samples generated full-length assemblies for 19 of the 24 genes in the respective tissue selected to sample for RNA-seq (Data File S1). Four of the remaining five genes could be assembled as either two (*WRKY8 and WRKY2)* or three (*HSF24* and *HSF20*) W22 transcripts due to small gaps in coverage and these genes have lower expression levels, except for HSF24 which has moderate expression, but only has 50 bp single-end sequencing which may contribute to the lack of complete assembly for this gene. The other gene, *HSF6*, lacked adequate coverage to assemble any mutant, *hsf6-m1* and *hsf6-m2*, or control transcripts and was removed from subsequent analyses. In total, transcript assemblies for 33 mutant alleles that aligned to the respective wild-type gene with a *Mu* insertion allele (23 genes) were identified and further characterized (Table S1, Data File S1). The transcripts from the mutant alleles were classified based on the presence of transcripts arising from the gene that are 5’ and 3’ of the *Mu* insertion or exclusively 5’ or 3’ of *Mu* (Figure 3A).

**Figure 3.**
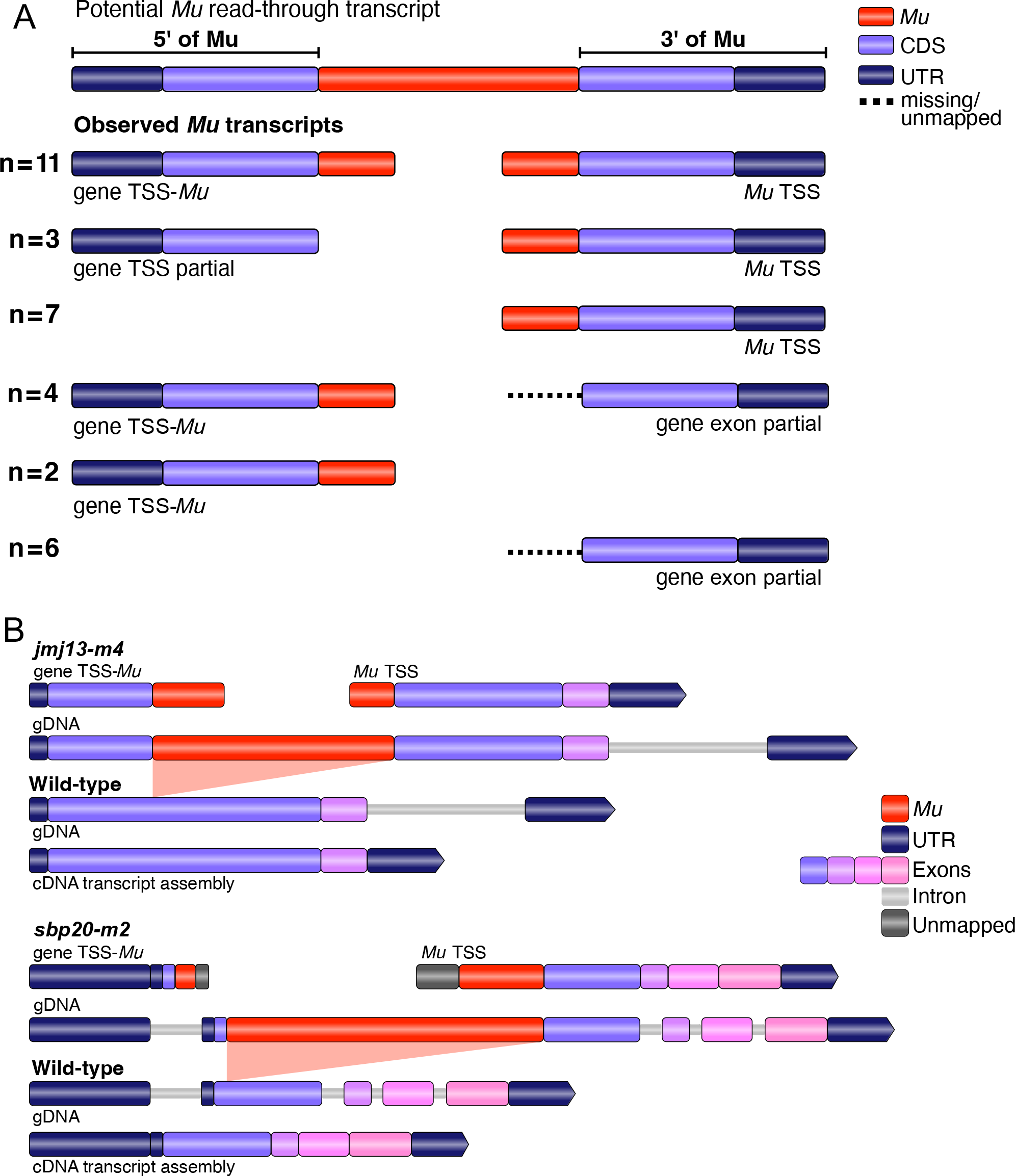
Schematic of *de novo* transcript assemblies for 33 mutant alleles. A) Mutant allele transcripts are referenced relative to the sequences 5’ and 3’ of the *Mu* insertion. No full- length transcripts with retained *Mu* sequences were identified. The observed transcripts could include separate transcripts representing gene sequences both 5’ and 3’ of the *Mu* insertion or sequences only 5’ or 3’ of *Mu*. We also identified some examples of transcripts with partial regions of retained *Mu* sequence. The observations were grouped into 6 transcript structure types that are illustrated with schematics and the total number of alleles for each type is indicated. Transcripts depicted as partial are truncated but contain no *Mu* sequence. **B)** The specific transcripts that are observed for two of the mutant alleles are shown in detail. The wild- type transcript assembly is shown to indicate we recovered the annotated wild-type cDNA with our short-read assembly. Both *jmj13-m4* and *sbp20-m2* have evidence for two transcripts assembled, one transcript initiating at the gene TSS with premature termination in *Mu* (gene TSS-*Mu*, 5’ of *Mu*) and the other initiating within *Mu* and reading through the 3’ end of the gene (*Mu* TSS, 3’ of *Mu*). Unmapped indicates assembled sequence that could not be annotated as *Mu* or gene sequence. All *de novo* transcripts were assembled with TRINITY.

For 20 alleles we identified an assembled transcript that matched gene sequences 5’ of the *Mu* insertion site (Figure 3A, Table S1, Data File S1). Many (17) of the 20 mutant alleles with assembled transcripts that contain gene sequence 5’ of *Mu* include at least a portion of *Mu* sequence at the 3’ end of the transcript which indicates the transcript reads into the *Mu* element and terminates. The 13 alleles (*Mu* TSS; n=7 and gene exon partial; n= 6, Fig. 3A) that did not have transcripts assembled 5’ of *Mu* are enriched for *Mu* insertions very near the 5’ end of the gene (*Mu* insertion sites within the first 33% of gene cDNA). In these cases, it is likely that if a short transcript (< 200 bp) was produced by the normal gene promoter it would be underrepresented in the RNA-seq data due to the size selection step during library preparation (Hirsch *et al*. 2015) and the lack of an assembly could reflect technical bias against short transcripts rather than absence of this transcript. The majority (31/33) of mutant alleles produce transcripts downstream (3’) of the *Mu* insertion site. Many (21/31) of these transcripts that include gene sequence downstream of the *Mu* insertion site contain a portion of recognizable *Mu* sequence at the 5’ end of the assembled transcript (Figure 3A, Table S1, Data File S1). The remaining 10 alleles have partial transcript coverage of gene sequence downstream of the *Mu* insertion and often (7/10 alleles) have relatively low expression levels (< 5.3 FPKM). Examples of assembled transcript structures observed for two *Mu* alleles, *jmj13-m4* and *sbp20-m2*, compared to the W22 allele are shown in Figure 3B. These mutant transcript assemblies suggest that many of the *Mu* insertions within the coding sequence result in the presence of two partial transcripts: a transcript initiating at the the gene TSS and terminating within *Mu* (gene TSS-*Mu*) or prematurely terminating (gene TSS partial) and a transcript initiating at a *Mu* TSS reading through the end of the gene (*Mu* TSS transcript).

### *Mu* promoter initiation does not strictly depend on a specific *Mu* element or orientation

There are multiple distinct members of the *Mutator* transposon family that could be mobilized, and insertions of these elements could occur in forward or reverse orientations relative to the gene sequence. To further understand the impact of specific *Mu* transposons and their orientation upon the potential of the *Mu* promoter to initiate outward-reading transcripts, we sought to characterize the identity and orientation of each *Mu* insertion mutant allele. The *Mu* sequence from each *de novo* assembled transcript, either present at the 3’ end of the gene TSS partial transcripts or at the 5’ end of the *Mu* TSS transcripts, was used to perform a BLAST search against representative examples of *Zea mays Mu* elements. *Mu* element identity was predicted based on the *Mu* element with the greatest similarity to *Mu* sequence from each transcript (Figure 4). Most (75%) of the assembled transcripts contain Mu sequences that align to the *Mu* terminal inverted repeats (TIRs) alone with only a subset that include internal *Mu* sequences (Figure 4). The transcript assembly *Mu* sequence alignments suggest that there are eight *Mu1* or *Mu1.7* elements (these cannot be separated based on the TIR regions alone), two *Mu3*, five *rcy:Mu7*, five *Mu8* and one *Mu13* element in our set of alleles. The 5’ and 3’ TIRs of *Mu* elements often have polymorphisms between the two TIRs and these wereused to predict the orientation of the *Mu* insertion for 15 of the 21 alleles with the presence of *Mu* sequence in their transcript assembly. One mutant allele with *Mu* inserted into an intron, *bzip76-m2*, had evidence for only internal *Mu* sequences, potentially suggesting that this allele may produce a *Mu* spliced transcript instead of a *Mu* TSS transcript. The predictions for *Mu* element identity and insertion orientation based on sequence alignments were tested using outward-reading PCR primers with specificity to either the 5’ or 3’ internal *Mu* sequence (Figure S2). We were able to confirm the identity for 14 of the 15 *Mu* insertions that had alignment-based predictions and for 12/14 of these the predicted orientation was supported by PCR-based testing (Figure S2, Table S3). The *Mu* element identity and orientation for the remaining alleles without discernible predictions for *Mu* orientation based on transcript alignments was determined using *Mu* element specific primers (Figure S2).

**Figure 4.**
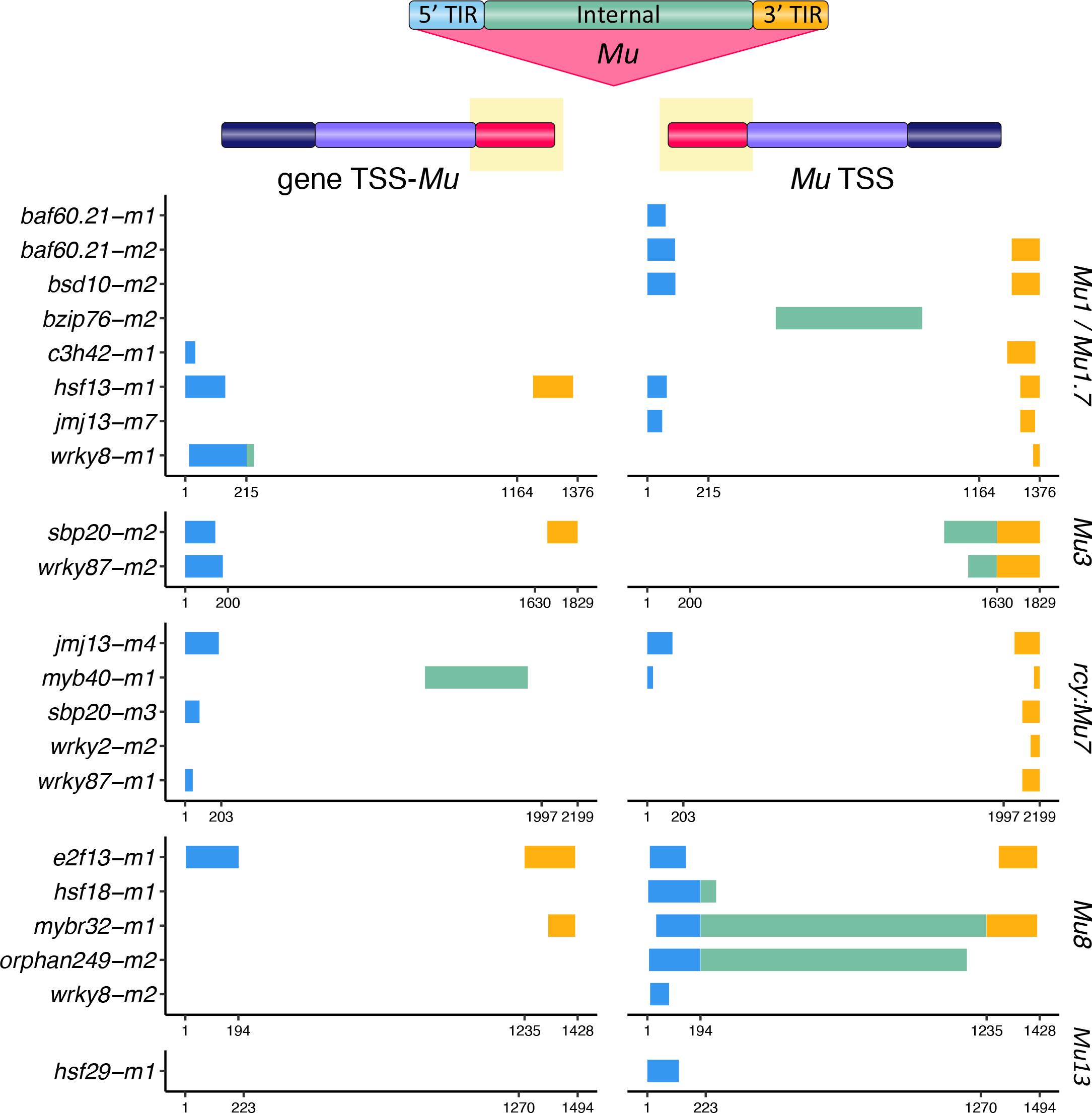
Assessment of *Mu* element identity and orientation. The *Mu* sequence from the *de novo* assembled transcripts (highlighted in red in the schematic) was aligned to representative *Mu* element sequences (element identity indicated to the right of the plots). The left panel shows alignments of the *Mu* sequence from the transcript initiated at the gene TSS while the right panel shows alignments of *Mu* sequence from *Mu* TSS initiated transcripts that include the 3’ portion of the gene. Mutant alleles are grouped by the *Mu* element with the greatest sequence similarity to the transcript assembled *Mu* sequence and the plots indicate the position of the aligned sequence within the *Mu* element. *Mu1* and *Mu1.7* are plotted together due to sequence similarity between TIR regions. The *Mu* sequence segments are colored by the alignment position relative to *Mu* element features: 5’ TIR (blue), internal sequence (green), and 3’ TIR (gold). The base pair coordinates listed in each plot indicate the TIR boundaries.

The observation that *Mu* elements in both forward and reverse orientations relative to the gene sequence appear to provide an outward-reading promoter suggested the potential for both bi- directional transcripts originating from the *Mu* element. The RNA-seq data was generated using strand-specific protocols. The analysis of read orientation between the control and mutant samples did not reveal excess antisense RNA-seq reads in the region 5’ of the *Mu* insertion (Figure S3). This suggests orientation-specific transcript initiation, potentially due to directional interactions with the endogenous promoter. Alternatively, it is possible that antisense transcripts are highly unstable and rapidly degraded and therefore not observed.

### Evidence for transcript termination and initiation within *Mu*

The observation that the majority of *Mu* insertion alleles within genes generate two separate transcripts could reflect the failure to assemble *de novo* transcripts through the *Mu* element or the presence of two independent transcripts, one initiating at the gene TSS and the other initiating at an outward-reading promoter within *Mu*. Our previous RT-PCR results (Figure 2C) suggest that at least a portion of *Mu* is not retained in a full-length transcript; however, these results could also reflect difficulty in successfully amplifying through the full *Mu* element (amplification of terminal inverted repeat sequences can be challenging). To rule out the potential of *Mu* insertion alleles producing transcripts that read through the *Mu* element, we performed RT-PCR using gene-specific primers and *Mu*-specific primers within and beyond the sequence that is observed in the *de novo* assembled transcripts (Figure 5A, Table S4). To ensure our RT-PCR *Mu* sequence amplification approach is comparable to the transcriptome assembly results, mutant and wild-type allele cDNAs were generated from the same tissue type sampled for RNA-seq. This approach allowed us to determine how much of the *Mu* element was retained in each mutant allele transcript tested.

**Figure 5.**
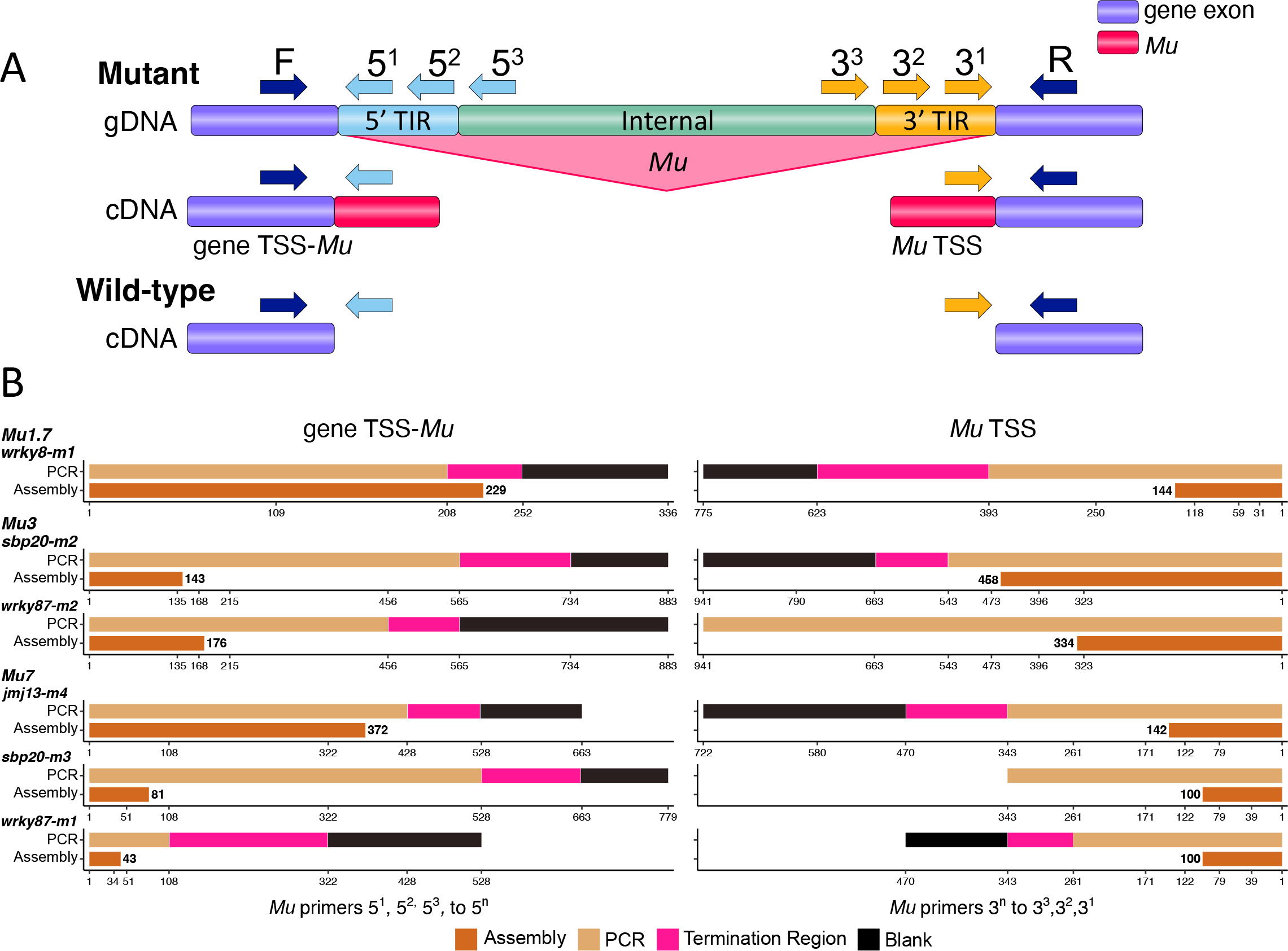
Definition of transcript boundaries using RT-PCR method. A) A schematic of how RT-PCR primers were designed to test the extent of the transcribed *Mu* sequence relative to the predictions of the mutant allele transcript assemblies. Multiple *Mu* primers were designed with specificity to either the 5’ or 3’ regions of each *Mu* element tested by RT-PCR. For each mutant allele, some *Mu* primers were designed to be located within the transcriptome assembly and some were designed for more internal regions of the *Mu* element. All primer-sets used for RT- PCR were first tested and confirmed to amplify mutant genomic DNA. **B)** A graphic representation of the RT-PCR results is shown. For each mutant allele, transcripts are categorized as gene TSS-*Mu* or *Mu* TSS with opposite orientations to mirror Figure 5A and alleles are ordered by Mu element identity. The numbers on the x-axis indicate *Mu* sequence length (bp) where each primer binds. Brown indicates *Mu* sequence regions expected to amplify based on the transcript assembly with the length of *Mu* sequence (bp) listed adjacent. Tan represents which primers successfully amplified products by RT-PCR. Pink indicates the first *Mu*-specific primer used that resulted in absence of amplification by RT-PCR and is the region where the transcript terminates. Black indicates absence of amplification by RT-PCR for *Mu* sequence that is more internal to the region where the transcript terminates (Blank). Absence or presence of amplification was confirmed by at least two biological replicates.

Transcript assemblies using RNA-seq data often fail to capture the full 5’ and 3’ ends of mRNA sequences. We noted that a number of the control assemblies lacked the full UTR sequence. In order to better assess the 3’ end of the gene TSS-*Mu* transcript and the 5’ end of the *Mu* TSS transcript, we assessed the presence of RT-PCR amplification products from a gene-specific primer and a primer with specificity to *Mu* sequence. The primers within *Mu* sequences included primers within the assembled sequences as well as several primers that were further into the *Mu* sequence. In all cases, we successfully amplified regions that were within the assembled transcripts (Figure 5B). The transcript boundaries were determined for 4/6 *Mu* TSS transcripts and all six of the gene TSS-*Mu* transcripts. For the two genes in which the *Mu* TSS initiated transcript boundary was not determined, there were issues in designing primers that amplified *Mu* internal sequence (*sbp20-m3*) or there was amplification of *Mu* sequence that was > 600 bp more internal than the assembly predicted (*wrky87-m2*). The majority of transcripts that were tested (11/12 for 6 alleles) resulted in successful amplification using primers that were slightly outside the assembled *Mu* sequence indicating that the assemblies are likely partially truncated. In most cases in which the transcript has identifiable boundaries (8/10), there were less than 300 bp of additional *Mu* sequence amplified. The other two transcripts reveal evidence for > 300 bp of additional *Mu* sequence beyond the transcript assembly. Together, these results confirm the lack of read-through transcripts but do suggest that the transcripts may read further into *Mu* or initiate further within *Mu* than revealed by the transcript assemblies (Figure 5B, Table S4).

### Transcripts from the mutant allele often have similar abundance to the wild-type allele

We were interested in comparing the expression level of each mutant transcript with the wild-type allele to document potential variability in transcript abundance. To compare expression levels between the mutant and wild-type alleles, we focused on RNA-seq reads that mapped to exon regions with shared sequence between the wild-type transcript and either the mutant gene TSS partial (terminated within *Mu* or prematurely terminated) or *Mu* TSS transcript (Figure 6A, Table S5). The expression level of shared exon regions (referred to as CPM per fragment) between the mutant gene TSS partial and wild-type transcripts reveal highly similar transcript abundances, R^2^ = 0.97 for 18 transcripts (Figure 6B, Figure S4). The *Mu* TSS transcript abundance was generally quite similar to the levels of the wild-type transcript, but *Mu* TSS transcripts exhibit more variation with an R^2^ = 0.43 for 19 transcripts (Figure 6C, Figure S4).

**Figure 6.**
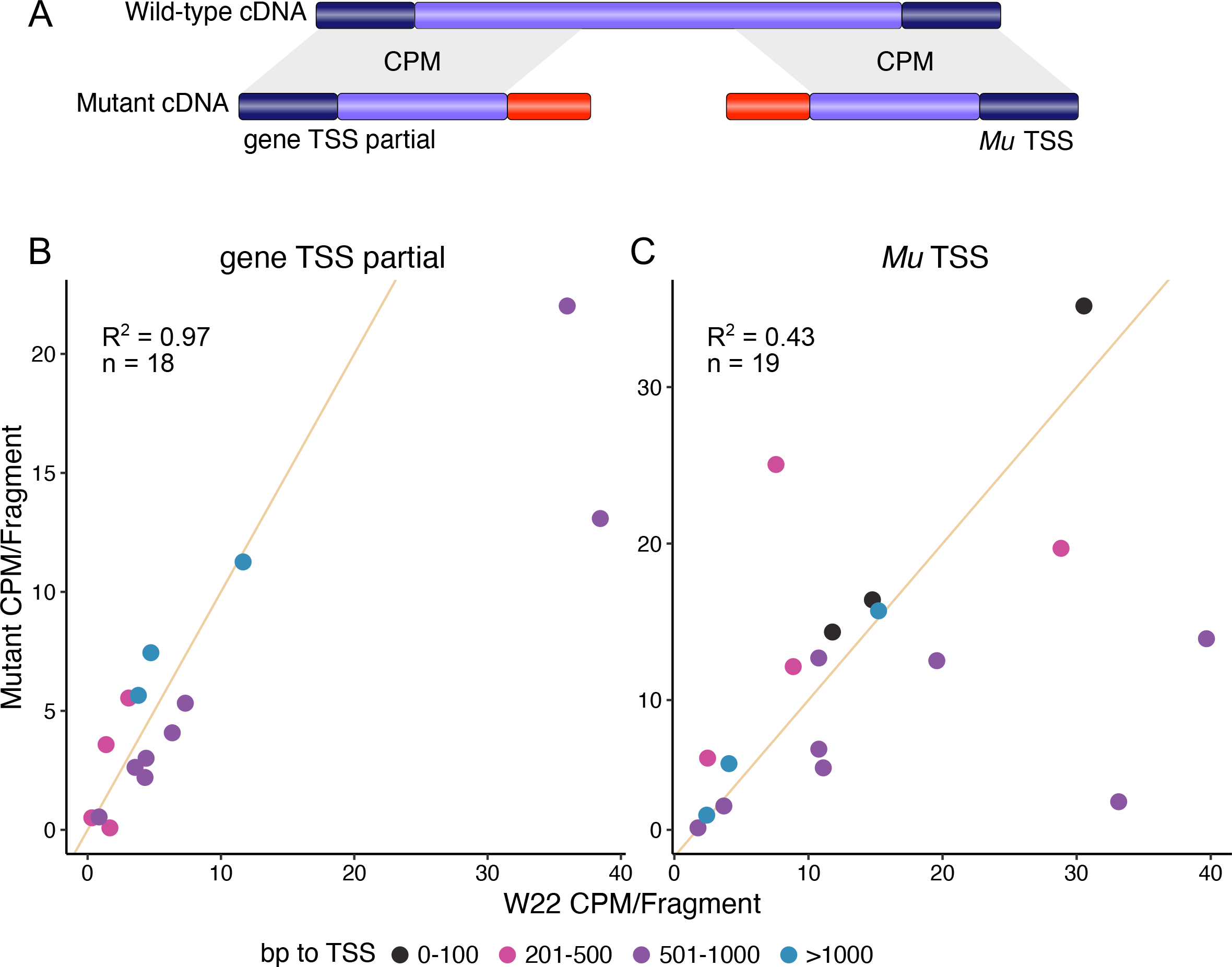
Transcript abundance comparison of mutant and wild-type transcripts. A) Schematic of how the expression level of the mutant and wild-type transcripts could be directly compared. Exon regions that are present in both the mutant and wild-type transcript assemblies were identified for each *Mu* allele. Transcript abundance (CPM/fragment) for mutant and wild- type transcripts was calculated from counts of the corresponding RNA-seq sample reads that map to these shared exon regions. A scatter plot of mutant transcript abundance (y-axis coordinates) relative to W22 wild-type transcript abundance (x-axis coordinates) for **B)** gene TSS partial transcripts, including transcripts with termination in *Mu,* from 18 mutant alleles and **C)** *Mu* TSS transcripts from 19 mutant alleles. Alleles are colored by the distance in bp of the *Mu* insertion from the gene annotated TSS: 0-100 (black), 201-500 (pink), 501-1000 (purple), > 1000 (blue). The lines show the expectation if there was equivalent expression for both alleles (slope = 1). Transcript abundance values for four mutant transcripts (*mybr32-m1*, *gras75-m1* and *wrky82-m1* gene TSS partial transcripts and *baf60.21-m1 Mu* TSS transcript) are not shown in these plots due to either mutant and/or wild-type abundance > 40 CPM. However, R^2^ values were calculated for each transcript abundance comparison from all 18 gene TSS partial transcripts and 19 *Mu* TSS transcripts.

There were several examples of lower expression levels for the *Mu* TSS transcript relative to the wild-type transcript (Figure 6C). The analysis of the distance of the *Mu* insertion site from the TSS did not suggest that the variation in expression levels for some *Mu* TSS transcripts was due to examples with more distant insertion sites. The observation that the *Mu* TSS transcript abundance is often similar to the wild-type in a single tissue suggests that this *Mu* outward-reading promoter may provide expression levels comparable to the wild-type gene promoter.

### *Mu*-derived transcripts can maintain similar tissue-specific patterns

We proceeded to assess whether mutant transcripts derived from the gene or *Mu* promoter would exhibit similar patterns of expression across multiple tissues. To evaluate tissue-specific expression, we performed RT-qPCR by amplifying transcript regions that are shared between the wild-type and mutant gene TSS-*Mu* or *Mu* TSS transcripts (Figure 7A, Table S6). To ensure we could test tissue-specific expression patterns, we selected genes with variable levels of wild-type gene expression across multiple tissues. Mutant gene TSS-*Mu* transcripts maintain relative expression levels that are very similar to wild-type transcripts across all tissues tested (Figure 7B) suggesting similar expression patterns for this mutant transcript and the wild-type transcript. The mutant *Mu* TSS transcripts often maintain wild-type tissue-specificity but frequently have lower relative transcript abundance–higher Delta Ct (Figure 7B). The *Mu* element identity (*Mu1.7, Mu3, rcy:Mu7* or *Mu8*) and *Mu* insertion position (5’ UTR or CDS) vary among the ten mutant alleles tested by RT-qPCR. The finding that all 10 mutant allele *Mu* TSS transcripts have patterns of expression similar to wild-type tissue-specificity suggests the ability of the *Mu* outward-reading promoter to mimic wild-type gene expression patterns is not entirely dependent on the specific *Mu* element or where *Mu* inserts within a gene.

**Figure 7.**
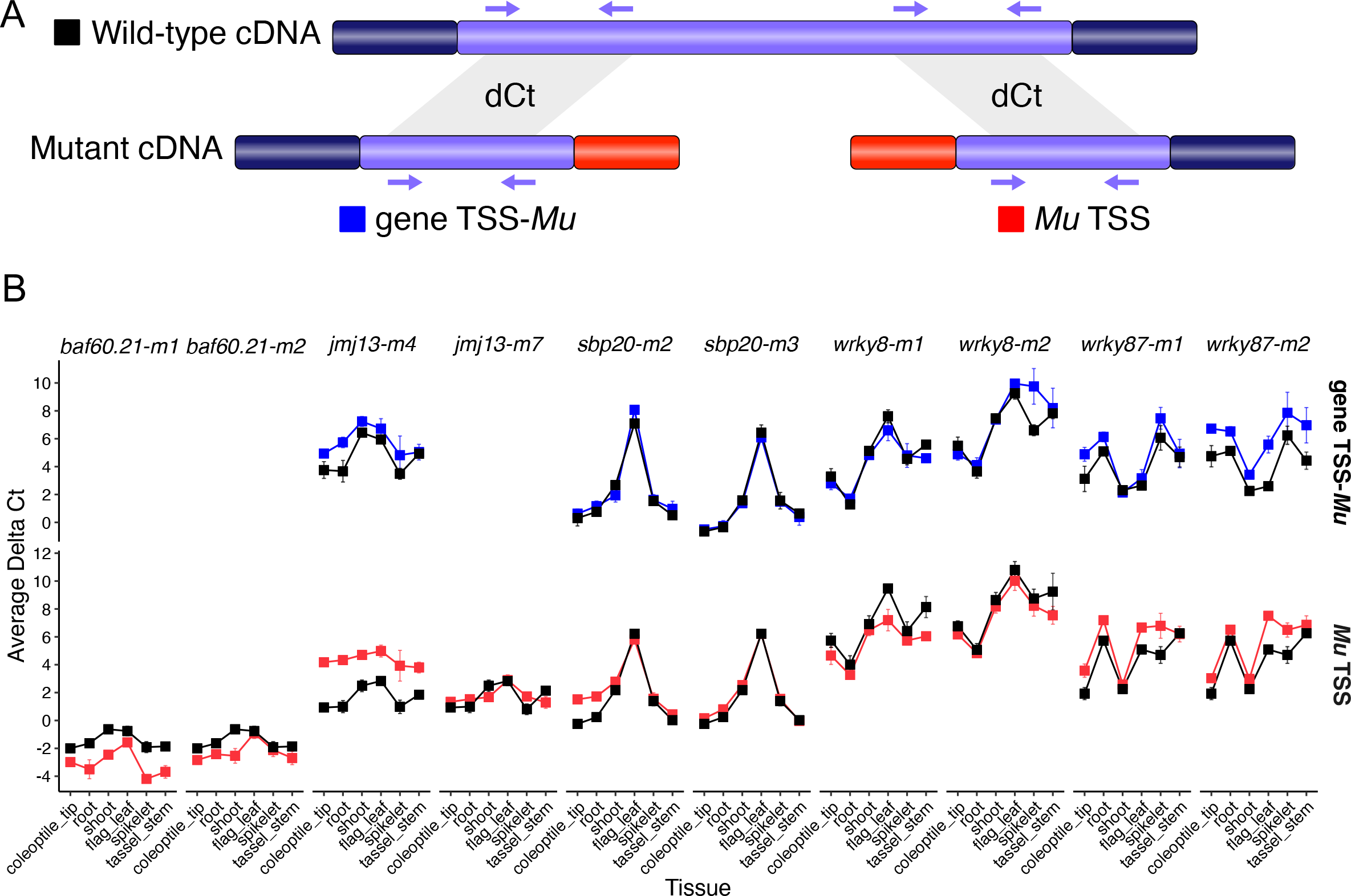
Tissue-specific patterns of expression for mutant transcripts relative wild-type transcripts from RT-qPCR data. A) Schematic of RT-qPCR primer design to test relative patterns of gene expression from mutant and wild-type alleles. Primers with specificity to regions of shared sequence between assembled wild-type and mutant transcripts were used to quantify tissue-specific expression patterns by RT-qPCR. 10 mutant alleles isolated from 5 genes were selected to sample for this analysis. Selecting genes with two independent mutant alleles each provided more power to confirm relative wild-type gene expression patterns across tissues and we focused on genes with variable expression levels in the tissues sampled. **B)** Average delta Ct (dCt) values of mutant gene TSS-*Mu* transcripts (blue) and mutant *Mu* TSS transcripts (red) relative to the values of their corresponding wild-type transcripts (black) are plotted as points each tissue sampled (x-axis tissue order is the same for all alleles). The gene TSS transcript for *wrky8-m2* is a gene TSS partial transcript. Each data point is the average dCt value of three biological replicates each with three technical replicates. Trendlines across tissues are plotted to display expression patterns. dCt values are inversely proportional to relative transcript abundance and greater dCt values indicate lower relative transcript abundance.

## Discussion

Previous work characterizing mutant alleles from insertions of *Mutator* transposons in maize genes identified the presence of an outward-reading promoter in *Mu (Barkan and Martienssen 1991; Chatterjee and Martin 1997)*. The ability of this *Mu* outward-reading promoter to initiate transcription is conditional upon the epigenetic state of *Mu (Barkan and Martienssen 1991)*.

*Mutator* elements can be in either one of two states: an active state (*Mu*-active) where there is a high forward mutation rate from the presence of active *MuDR* transposons or an inactive state (*Mu*-inactive) without *MuDR* activity (Chomet *et al*. 1991b). The state of *Mu* activity can be monitored by the extent of DNA methylation in sequences of *Mu* terminal inverted repeats (TIRs); plants with *Mu*-active state elements are marked by hypomethylation and plants with*u*-inactive exhibit hypermethylation (Bennetzen 1984; Chandler and Walbot 1986; Barkan and Martienssen 1991; Chandler and Hardeman 1992; Raizada and Walbot 2000). There are several reports demonstrating that insertions of *Mu* transposons into maize genes can lead to mutations whose phenotypes are suppressed only in the absence of *Mu* activity (*Mu*-inactive) (Martienssen *et al*. 1990; Chomet *et al*. 1991b; Lowe *et al*. 1992; Greene *et al*. 1994; Das and Martienssen 1995; Hu *et al*. 1998; Girard and Freeling 2000; Settles *et al*. 2001; Cui *et al*. 2003). In the *Mu*-inactive state, the *Mu* promoter becomes active and initiates transcription directed outward into the adjacent gene to restore the phenotype of the *Mu*-induced allele to that of its progenitor (Barkan and Martienssen 1991). Mutant alleles with phenotypes that depend on the activity of *Mu* for expression are known as *Mu*-suppressible alleles (Martienssen *et al*. 1989).

In our study, we provide evidence that transcript initiation in *Mu* and potential activity of a *Mu* outward-reading promoter is a common phenomenon for mutant alleles isolated from the UniformMu mutant population in maize. Most mutant alleles (20/33) isolated in our study have transcript assembly evidence of transcript initiation in *Mu* sequence. Although we did not directly examine *Mu* activity in these stocks, we can infer that these mutant alleles are likely in a *Mu*- inactive state as one step in the creation of the UniformMu population includes selection for kernels that lack evidence of *Mu* activity prior to identification of insertion alleles (McCarty *et al*. 2005). Previously reported *Mu*-suppressible alleles in maize with evidence of an active promoter in *Mu* were generated from a *Mu* insertion in the promoter or 5’ UTR of a gene (Barkan and Martienssen 1991; Chatterjee and Martin 1997; Girard and Freeling 2000; Settles *et al*. 2001; Pooma *et al*. 2002; Cui *et al*. 2003). However, the previously characterized *Mu*-suppressible alleles were all detected based on phenotypic effects and likely required production of transcripts from the *Mu* promoter that could produce a functional protein. *Mu* elements inserted near the wild-type TSS provide the potential for production of a transcript that can encode the full ORF of the gene. However, these insertions near the 5’ end of the gene may also allow for activity of the outward-reading promoter within *Mu* based upon position of this *Mu* promoter very near the site of the gene promoter which may allow for the *cis*-regulatory elements of the gene to influence this *Mu* promoter.

Our results show that the activity of the outward-reading promoter in *Mu* is not dependent on the *Mu* insertion site being in the gene promoter or 5’ UTR. For the 20 alleles with evidence of a *Mu* promoter initiated transcript, 13 were isolated in coding sequences from various positions spanning the gene length, five were within the 5’ UTR, and two were within introns. These results suggest that the distance of the *Mu* promoter from the wild-type gene promoter does not necessarily determine activity of the *Mu* promoter. Previous reports indicate the *Mu* outward-reading promoter is located near the edge of the *Mu* element–potentially initiating transcripts from within the *Mu* TIR sequence (Barkan and Martienssen 1991; Chatterjee and Martin 1997). When we aligned sequences from *Mu* TSS transcripts of 20 alleles to predicted *Mu* element sequences, the *Mu* transcribed sequence from 15/20 transcripts mapped entirely to the TIR sequence. Although the exact location of the promoter within *Mu* was not precisely defined, the *Mu* outward-reading promoter is likely located near the termini of *Mu*. We find examples of multiple Mu elements (*Mu1.7, Mu3, rcy:Mu7* and *Mu8*) and *Mu* insertions in either the forward or reverse orientation relative to the gene TSS can provide an outward-reading promoter. Several prior studies also suggested that *Mu*-suppressible alleles could include different types of *Mu* elements and orientations (Greene *et al*. 1994; Girard and Freeling 2000)

The UniformMu induced mutations are a widely used tool for functional genomics in maize (McCarty *et al*. 2005, 2013, 2018; Settles *et al*. 2007; Liu *et al*. 2016). While the silencing of *Mu* transposition is quite useful for ensuring that the detected *Mu* insertions represent germinal rather than somatic insertion events, it also has the potential to create *Mu*-suppressible alleles through potential activation of *Mu* outward-reading promoters. Our study provides evidence that for many UniformMu mutant alleles the *Mu* element provides an outward-reading promoter that can direct transcription into adjacent gene sequences. *Mu* promoter activity seems to be similar to *Mu* suppression in terms of frequency, as it has been reported that in the absence of *Mu* activity over half of the *Mu*-induced mutations are suppressed (May *et al*. 2003). Although we found that *Mu* promoter activity does not depend on insertion site, it is known that *Mu* preferentially inserts into promoter and 5’ UTR regions (Dietrich *et al*. 2002; Vollbrecht *et al*. 2010; Springer *et al*. 2018). This means that many *Mu* insertion alleles within the UniformMu population represent 5’ UTR insertion events. Researchers should use caution when interpreting the phenotypic results of these insertion events as it is possible that *Mu* promoter initiated transcripts could complement the insertion. Failure to obtain RT-PCR products when using primers that flank the *Mu* insertion site does not necessarily indicate a true loss-of-function allele as a *Mu* initiated transcript could still be produced. *Mu* insertions into regions upstream of the gene coding sequence are not only the most abundant (42%) in the UniformMu population (McCarty *et al*. 2013; Andorf *et al*. 2016), but also allow the potential for the promoter in *Mu* to drive a transcript that includes the entire original ORF and complement the mutant phenotype.

The tissue-specificity of the *Mu* outward-reading promoter has not been well characterized. The ability of the *Mu* outward-reading promoter to suppress phenotypes of *Mu* insertion alleles suggests the ability to drive expression in tissues in which the gene product is needed to normally function. Prior work on *hcf106-mum1* suppressible alleles revealed that both the wild-type *Hcf106* and *hcf106:Mu* exhibit similar patterns of expression response to light (Barkan and Martienssen 1991), albeit with higher levels of expression for the wild-type gene compared to *hcf106:Mu*. Our collection of transcripts initiated within *Mu* provided an opportunity for a broader characterization of the activity of the *Mu* outward-reading promoter. Our findings suggest that the outward-reading promoter seems to be able to mimic the gene promoter in terms of expression level and tissue-specificity and has limited inherent patterns of expression. The RNA-seq data was generated from multiple tissues depending on which tissues exhibit high levels of expression for the wild-type gene. We found that the transcript abundance (CPM per fragment) of the mutant *Mu* TSS transcripts tend to be similar to wild-type for most alleles and that there were not consistent differences in the expression of the *Mu* TSS transcripts among the tissues. Several genes that were selected for characterization of expression patterns by RT-qPCR normally exhibit variation in expression among the profiled tissues. We found that the *Mu* TSS transcripts tended to mimic these tissue-specific patterns. Although *Mu* TSS transcripts follow wild-type tissue-specific patterns, they often exhibit expression levels that are slightly lower than that of gene TSS partial transcripts relative to wild-type. Our results, along with previous reports (Chomet *et al*. 1991a; Barkan and Martienssen 1991; Lowe *et al*. 1992; Settles *et al*. 2001; Cui *et al*. 2003), imply that *Mu* might provide a minimal outward-reading promoter that can interact with genic *cis*-regulatory elements to condition expression patterns and levels that are similar to the wild-type gene.

There are several mechanisms by which maize transposons can minimize the functional impact of insertions into genic regions. The *Mu* family of transposons seems to have adopted a mechanism of providing an outward-reading promoter that is active when the *Mu* is silenced.

The coupling of this mechanism with the preferential insertion within promoters and 5’ UTRs provides the opportunity for *Mu* elements to insert within open chromatin regions while limiting potential deleterious consequences. Our findings and previous studies (Barkan and Martienssen 1991) suggest that the *Mu* promoter relies on interactions with genic *cis*-regulatory elements to mimic wild-type gene expression patterns. By providing a minimal promoter that can mimic gene expression patterns *Mu* elements have the ability to insert and increase in copy number with limited effects on the long-term survival in the host. This provides an elegant solution for a transposon to potentially limit the consequences of its proliferation.

This study provides evidence that *Mu* transposon insertions often result in complex transcripts for the gene rather than clear loss-of-function alleles. Our mutant allele transcript assemblies frequently include examples of termination and initiation in *Mu* sequence. Transcripts initiating from *Mu* are likely derived from a *Mu* outward-reading promoter that may produce functional transcripts if these include the full ORF of the gene. These results have implications for the many researchers that utilize UniformMu for reverse genetics. Further studies are necessary to document whether the *Mu* outward-reading promoter requires a *Mu*-inactive state and to uncover the mechanisms that allow the *Mu* promoter to interact with genic *cis*-regulatory elements. The ability of *Mu* to provide an outward-reading promoter also has implications for future transposon biology. The system by which conditional activity of a *Mu* promoter determines whether *Mu* can suppress a mutant allele should be utilized to understand the relationship between transposons and host genomes.

## Methods

### Isolation of homozygous mutant alleles from the UniformMu population in maize

Transposon-indexed seed stocks were ordered from MaizeGDB Stock Center (Lawrence *et al*. 2004; McCarty *et al*. 2005). Most alleles were selected based on insertions into the coding sequence or 5’ UTR. Seeds were planted in the field to maintain seed stocks. At the Vegetative 3 (V3) developmental stage leaf tissue was collected for DNA isolation. Mutant alleles were genotyped to identify the presence and zygosity of *Mu* with gene-specific primers flanking the *Mu* insertion and a primer with specificity to the *Mu* TIR regions: 9242, as described in (McCarty *et al*. 2013). Homozygous transmissible alleles were then isolated after backcrossing twice to the W22 r-g inbred if possible, to reduce the original mutation load from the transposon-indexed stock (Table S1).

### Plant material for RNA-seq samples

Wild type tissue-specific expression data from B73v4 (Zhou *et al*. 2019) and W22 (Monnahan *et al*. 2020) were used to identify tissues where each of the 24 maize genes with a *Mu*-insertion mutant allele had moderate to high expression. To capture expression of each of the 24 genes in both mutant and control conditions, 5 different tissues were selected to sample: coleoptile tip, seedling leaf, imbibed embryo, tassel, and tassel stem (Table S2). Three biological replicates of the mutant allele and at least three biological replicates of control W22 r-g were sampled for RNA-seq from the respective tissue selected for each gene (Table S1). Samples for each tissue were collected on the same day at the same time with the 24 h time sampled listed in parentheses. At anthesis, tassel stems (∼3 cm, anthers extruded, 9:00) and whole tassels (anthers unextruded, 12:00) were sampled from three plants in the field each and pooled for one biological replicate. Embryos were dissected (from 8:00-11:00) after imbibing seeds in distilled water for 48 h at 31°C and 5 embryos were pooled for one biological replicate. For seedling leaf tissue, the V3 collared leaf was sampled (9:00) from each seedling 10 days after sowing (DAS) in 16 h light 28°C, 8 h dark 24°C growth chamber conditions and 3 leaves were pooled for each biological replicate. Coleoptile tips were sampled (9:00) from seeds 6 DAS in 30°C dark conditions using a paper towel cigar roll method for germination (Zhu *et al*. 2005) and 3 tips (∼2.5 cm) were pooled for each biological replicate.

### RNA-seq data processing

Total RNA was extracted using the RNeasy Plant Mini Kit (QIAGEN, Cat # 74904), quantified internally and externally by University of Minnesota Genomics Center (UMGC) with the Quant-iT RiboGreen RNA Assay Kit (Thermo Fisher, Cat # R11490), and quality checked with the Agilent 2100 Bioanalyzer. One biological replicate of the mutant allele, *gras52-m1*, had low RNA quality and was discarded prior to sequencing. Sequence libraries were prepared from a minimum of 500 ng of total RNA using the standard TruSeq Stranded mRNA library protocol (Illumina, Cat # 20020595) and sequenced on the NovaSeq 6000 S4 flow cell to produce at least 20 million 150 bp paired-end reads for each sample. For all samples with paired-end sequencing, both library construction and sequencing were done at UMGC. Library construction and sequencing for two mutant alleles, *hsf24-m3* and *hsf24-m4*, and W22 control sampled from tassel tissue was done externally at the Genomic Core at Michigan State University. For these samples, libraries were prepared from 2 µg of total RNA using the TruSeq RNA Sample Prep Kit (Illumina, Cat # FC- 122-1001) and sequenced on the HiSeq 4000 to produce at least 18 million 50 bp single-end reads.

For all samples, sequencing reads were then processed through the nf-core RNA-Seq pipeline (Di Tommaso *et al*. 2017; Ewels *et al*. 2020) built with Nextflow v20.10.0 (Di Tommaso *et al*. 2017) for initial QC and raw read counting. Reads were trimmed using Trim Galore! v0.6.5 (Krueger) and aligned to the W22 reference genome (Springer *et al*. 2018) using Hisat2 v2.1.0 (Kim *et al*. 2015) with default parameters (“hisat2 -x $db $input -p 12 --met-stderr --new-summary”). Uniquely aligned reads were counted per feature by featureCounts v2.0.1 (Liao *et al*. 2014). Raw read counts were normalized by library size and corrected for library composition bias using the TMM normalization approach in edgeR v3.28.0 (Oshlack *et al*. 2010), to give CPMs (Counts Per Million reads) for each gene in each sample allowing direct comparison between mutant and control samples (Table S1). CPM values were normalized by gene CDS lengths to give FPKM (Fragments Per Kilobase of exon per Million reads) values (Table S1). Genes were considered expressed if their CPM was ≥ 1 in at least one sample per tissue.

### Identification of differentially expressed genes

Raw read counts of expressed genes (CPM ≥ 1 in at least 1 sample per tissue) from all replicates of each mutant allele and W22 control from the same tissue were used to call differentially expressed (DE) genes, false discovery rate [FDR] adjusted p-value < 0.05 and a minimum fold change of 2 (DESeq2 v1.30.1 (Love *et al*. 2014)) (Table S1).

### Transcriptome profiling

Reads from RNA-seq data of combined biological replicates for each allele, mutant or control, were trimmed with Trimmomatic v0.33 (Bolger *et al*. 2014) and *de novo* assembled into transcripts with TRINITY v2.5.1 (Grabherr *et al*. 2011) using default parameters. A local blast database (SequenceServer (Priyam *et al*. 2019)) was created for each *de novo* transcriptome assembly to identify transcripts aligning to the W22 gene cDNA in both the mutant and control. W22 control transcript assemblies for each gene were analyzed first by both BLASTn (Altschul *et al*. 1990) and the ExPASy translate tool (Duvaud *et al*. 2021) to confirm TRINITY could assemble the full-length gene cDNA from the RNA-seq short-read data. The canonical ORF of each gene was identified by comparing the annotated W22 gene cDNA sequence to sequences of orthologous genes in other grass species (i.e., *Sorghum Bicolor, Setaria Italica, Oryza sativa*) via BLASTx. To determine if Mu sequence was transcribed, the transcript assembly sequence of each mutant allele was searched against all standard nucleotide databases by BLASTn without specifying an organism (Altschul *et al*. 1990). The effect of the transposon insertion for each mutant allele was predicted by examining the putative ORF. A six-frame translation was completed for each mutant and wild-type transcript with the Expasy translate tool (Duvaud *et al*. 2021) to identify the mutant ORF with shared sequence to the corresponding wild-type transcript and detect upstream ORFs within Mu sequence (Table S1, Data File S1).

### Predictions of *Mu* element identity and orientation

Sequence from each mutant assembled transcript was used as a query against all public sequencing databases–NCBI to identify if there were any hits to *Mu* elements (Data File S1). Complete sequences of representative *Zea mays Mu* elements with transcript assembly hits: *Mu1* (X00913.1), *Mu1.7* (Y00603.1), *Mu3* (JX843286.1:132-1963), *Mu4* (X14224.1), *Mu5* (X14225.1), *rcy:Mu7* (X15872.1), *Mu8* (X53604.1), Mu13 (HQ698272.1), *Mu17* (HQ698276.1), and *MuDR*-*MudrA* and *MudrB* (M76978.1), were used to create a local *Mu* element BLAST database (SequenceServer (Priyam *et al*. 2019)) (GenBank Nucleotide Accessions from NCBI). *Mu* sequence from each assembled transcript was then BLAST against only *Mu* element sequence and top hits were used to predict the *Mu* element for each allele (Data File S1). The *Mu* sequence from assembled transcripts of each mutant allele was then aligned to the complete sequence of the predicted *Mu* element. *Mu* element insertion orientation could be predicted when transcribed *Mu* sequence either only aligned to or aligned with greater similarity to 5’ or 3’ regions of *Mu*.

### PCR confirmation of *Mu* element identity and orientation

For each predicted *Mu* element, outward-reading primers with specificity to either the 5’ or 3’ sequence of *Mu* were designed (Table S3). We refer to a forward orientation of *Mu* when *Mu* 5’ TIR sequence relative to 3’ TIR sequence is closest to the gene TSS and reverse orientation when Mu 3’ TIR sequence is closest to the gene TSS. PCR was performed on mutant allele gDNA with specific combinations of gene-specific (referred to as F and R) and *Mu*-specific primers (referred to as 5 and 3) to confirm the identity and orientation of *Mu* (Table S3). The presence of amplicons from F:5 and/or R:3 indicates a *Mu* element with forward orientation while amplification from F:3 and/or R:5 indicates reverse orientation. The *Mu* primers designed had specificity to 5’ or 3’ sequences of a specific *Mu* element. Amplification of gDNA using these *Mu* element-specific primers was considered *Mu* element identity confirmation.

### Mutant assembled transcript structure assessed by RT-PCR

*Mu* sequences from transcript assemblies of all mutant alleles with a shared *Mu* element identity were aligned to the complete sequence of that *Mu* element. Outward-reading PCR primers were designed with specificity to regions of the *Mu* sequence included in the transcript assembly and regions outside of the assembly for each mutant allele (Table S4). Gene-specific primers flanking the *Mu* insertion were designed with specificity to both mutant allele gDNA and cDNA sequence (Table S4). For each mutant allele, PCR was performed on both gDNA and cDNA to determine if mutant transcripts terminated and initiated in *Mu* at the predicted transcript assembly sites. As a control, PCR was first performed on mutant allele gDNA to detect presence of amplification from each primer-set designed with specificity to gene and *Mu* sequence included in and outside of the transcript assembly. Combinations of gene-specific and *Mu*-specific primers that amplified mutant gDNA were then used to test for presence of amplification from mutant cDNA (Table S4). To test the predicted transcript structures for six mutant alleles by RT-PCR, total RNA was extracted from the same tissue type sampled for RNA-seq. Extracting RNA from the same tissue type RNA-seq was performed on allows for direct comparison between our RT-PCR results and the assembly results without bias of amplification from tissue-specific isoforms. Tissue was sampled for at least three biological replicates of each mutant allele and W22 control using tissue sampling methods listed above in the Plant Material section. Total RNA was extracted from ∼100 mg of tissue/sample using TRIzol^™^ Reagent (Thermo Fisher, Cat # 15596026), DNase treated (TURBO DNA-free™ Kit, Thermo Fisher, Cat # AM1907), and quantified with Quant-iT RiboGreen RNA Assay Kit (Thermo Fisher, Cat # R11490). RT-PCR reactions for each primer set were performed on RNA (50-75 ng/ul) from at least two biological replicates of the mutant allele and control (QIAGEN OneStep RT-PCR Kit, QIAGEN Inc., Cat # 210212). The bounds of where transcriptional termination and initiation occurs within *Mu* for each mutant allele was determined by presence of amplification from gDNA and absence of amplification from cDNA. If there was amplification of mutant cDNA from primers designed to amplify regions outside of the transcript assembly, then more *Mu* sequence was transcribed than the assembly predicted.

### Transcript abundance (CPM per fragment) calculations

Mutant assembled transcripts were aligned to wild-type assembled transcripts via MAFFT v7 (Katoh and Standley 2013) in Benchling and regions of shared sequence were identified.

Genomic coordinates of these shared exon regions were used to create an annotation file (BED format) for each gene. BAM files (from uniquely aligned RNA-seq reads previously mapped to the W22 genome) for each of the three biological replicates of mutant and W22 control were converted to BED format (BEDTools bamtobed (Quinlan and Hall 2010)). The RNA-seq mapped read BED file of each mutant or control biological replicate was intersected with the shared exon read annotation file to obtain new read counts (BEDTools intersect-force strandedness (Quinlan and Hall 2010)). Read counts were then normalized by the effective library size with edgeR v3.28.0 (Oshlack *et al*. 2010) to give CPMs (Counts Per Million reads) for each gene in each sample. Transcript abundance was calculated by averaging the CPMs from all three biological replicates of each mutant allele or W22 control (Table S5). The coordinates of shared sequence regions between mutant and wild-type transcripts usually span multiple exons; therefore, we refer to this transcript abundance calculation as CPM per fragment. Linear regression was used to calculate R^2^ correlation values between mutant and wild-type transcript abundance (lm function in R (R Core Team 2020)).

### Tissue-specific expression patterns of mutant transcripts

Five genes (two independent mutant alleles each) with variable levels of wild-type gene expression across multiple tissues were selected to analyze by RT-qPCR. To examine tissue-specific patterns of expression for each of the 10 mutant alleles relative to wild-type, 7 different tissues at various stages of maize development (immature to mature) were selected to sample: imbibed embryo, coleoptile tip, seedling shoot, seedling radical root, flag leaf, tassel stem, and immature ear spikelet. Three biological replicates of each mutant allele and W22 control were sampled for RNA from each of the seven tissues. The same sampling methods listed above in the Plant Material section were used to sample imbibed embryo, coleoptile tip, and tassel stem tissues. Coleoptile tips and radical roots were sampled (9:00) from the same plants, from seeds 6 DAS in 30C dark conditions, and 6 tips and roots were pooled for each biological replicate.

Whole shoots from three seedlings 10 DAS in 16 h light 28°C, 8 h dark 24°C growth chamber conditions were sampled (9:00) and pooled for one biological replicate. Along with tassel stems (11:30), unfertilized ear spikelets and flag leaves were sampled (10:30) at anthesis and tissue from three plants was pooled for one biological replicate. Unfertilized ear spikelets were sampled by trimming the terminal 4 cm of the ear and collecting the next 2 cm. For flag leaves, the terminal 15 cm of the flag leaf blade was collected.

Total RNA was extracted from ∼100 mg of tissue per sample using TRIzol^™^ Reagent (Thermo Fisher, Cat # 15596026), DNase treated (TURBO DNA-free^™^ Kit, Thermo Fisher, Cat # AM1907), and quantified with Quant-iT RiboGreen RNA Assay Kit (Thermo Fisher, Cat # R11490). RNA from other tissues was diluted to 75 ng/ul prior to RT-qPCR. Primers for RT-qPCR were designed with specificity to amplify shared gene sequence (∼300 bp) between mutant and wild-type transcript assemblies: regions 5’ of *Mu* sequence in mutant gene TSS-*Mu* transcripts and regions 3’ of *Mu* sequence in mutant *Mu* TSS transcripts (Table S6). The Luna^®^ Universal One-Step RT-qPCR Kit (New England Biolabs, Cat # E3005X) and reaction protocol (cycle variations: initial denaturation for 2 min., 39 cycles of denaturation/extension, and melt curve 65C-95C at 0.5C increments) was used to run all RT-qPCR reactions. For each primer-set and tissue, RT-qPCR was performed on three technical replicates of each biological replicate for three biological replicates of the mutant allele and control. Technical replicate Ct values were averaged for each biological replicate. Delta Ct (dCt) values were calculated by the difference between the gene Ct value and the Ct value of the selected maize housekeeping gene, *Ubiquitin Carrier Protein* (*Zm00004b005988*) (Manoli *et al*. 2012) for each biological replicate. Then, dCt values for all three biological replicates were averaged to give a final dCt value for each transcript (Table S6).

### Data and code availability

All datasets and scripts necessary for data analysis are available in the GitHub repository at https://github.com/erikamag/Mu-Pro_manuscript. The RNA-seq data generated for this study is available and NCBI SRA PRJNA936808.

## Acknowledgements

We thank the Minnesota Supercomputing Institute at the University of Minnesota (http://www.msi.umn.edu) for providing resources that contributed to the research results reported within this article. We thank Chase Dickson, Hayden Hamsher and Noelle Lynch for helping with data generation.

---

Figure S1. Visualization of wild-type W22 and mutant allele RNA-seq read coverage for three genes.

Figure S2. Determination of Mu element identity and orientation by PCR using Mu element specific primers.

Figure S3. Analysis of RNA-seq read orientation between wild-type W22 and the mutant allele for two genes.

Figure S4. Ratios of mutant to wild-type transcript abundance.

Table S1. 35 Mutator mutant alleles isolated from the UniformMu population in maize.

Table S2. Gene expression values for 24 transcription factor genes in different tissues.

Table S3. Mutant allele Mu element identity and orientation by gDNA PCR.

Table S4. Mutant allele transcript boundaries and potential for Mu read-through tested by RT-PCR.

Table S5. Transcript abundance for shared exon sequence between mutant and wild type transcripts.

Table S6. Tissue-specific expression patterns for mutant and wild type W22 transcripts tested by RT-qPCR.

